# Post-fire changes in plant growth form composition in Andean páramo grassland

**DOI:** 10.1101/2020.04.25.061036

**Authors:** Maya A. Zomer, Paul M. Ramsay

## Abstract

**Questions:** Fire suppression policies have been widely adopted in the páramo grasslands of the northern Andes to protect their biodiversity and ecosystem services. Páramos have been regularly burned for many years, and it is not clear how páramo vegetation will respond to significant changes in their fire regimes. This study investigates differences in plant growth form composition, light levels and soil temperatures in páramo plots representing a range of recovery times since the last fire.

**Location:** Reserva Ecológica El Ángel and La Bretaña Nature Reserve, Carchi, Northern Ecuador.

**Methods:** We assessed the frequency of ten páramo growth forms, vegetation height, soil temperature, and light intensity in fifteen fire sites with historical records of fire (<1 – 15 years since fire), and one recently unburned site (at least 40 years since fire). A chronosquence of sites was used to assess potential changes in plant community composition in post-fire succession of páramo.

**Results:** The recovery of páramo vegetation after fire comprised three phases: initial recruitment with high growth form diversity, followed by reduced diversity, light and soil temperatures in dense tussock vegetation, and ultimately canopy height stratification with a return of diversity. All but one plant growth forms were represented in each of the three phases, and the changes reflected differences in relative abundance.

**Conclusions:** Post-fire páramo succession is characterized by clear shifts in the relative abundance of plant growth forms, ending with (co-)dominance of upright shrubs. The long-term consequences of such shifts for biodiversity and ecosystem function, given the widespread adoption of fire suppression policies in the páramo need careful, evidence-based consideration.

## Introduction

The páramos are the largest extension of tropical alpine ecosystems, forming a discontinuous belt throughout the northern Andes, with outliers in Panama and Costa Rica. The páramo has the most diverse mountain flora in the world (Smith and Cleef 1988), and these grasslands sustain ecological processes, carbon storage, and supply water for millions of people, agriculture and industry at lower latitudes (Buytaert et al. 2011).

Fires are the most significant human impact in the páramos (Horn and Kappelle 2009; Laegaard 1992; Ramsay and Oxley 1996) and people have burned these grasslands for thousands of years (White 2013). Putative fire suppression policies have been introduced to counter a reported increase in burning frequencies in some areas (Armenteras et al. 2020), a conservation strategy that stems from the common perception of fire as a threat to ecosystem integrity and services in the high Andes (Horn and Kappelle 2009; Keating 2007; Matson and Bart 2013). Understanding biodiversity and community level responses to such strategic decisions in the páramo requires an in depth knowledge of both fire regimes and how páramo vegetation recovers through time after fire (Matson and Bart 2013; Ramsay 2001).

Fire disturbance might represent a powerful mechanism of promoting and maintaining species diversity in páramo grasslands (Horn and Kappelle 2009; Keating 2007; Sklenář and Ramsay 2001). Regular burning in the páramo promotes a destruction-renewal cycle that has resulted in a landscape composed of a mosaic of patches in varying stages of recovery (Grubb 1977; Ramsay 1999; Ramsay 2001; Smith and Young 1987). Fine-scale heterogeneity in vegetation structure leads to patchy distribution of fuel and variable fire temperatures, which results in differential plant mortality, and post-fire establishment and growth (Ramsay 2001; Ramsay and Oxley 1996). Such patchiness and variability in recovery after fire have been linked to higher levels of biodiversity at the landscape scale (Keating 2007; Sklenář and Ramsay 2001) and at a finer scales (Ramsay and Oxley 1996; Sarmiento and Frolich 2002).

Studies of post-fire vegetation development have been carried out in Ecuador, Colombia, Venezuela, and Costa Rica and have shown variable changes in composition, cover and stature (Horn and Kappelle 2009). Only a few studies have monitored vegetation recovery through time after fires in Ecuador. Bremer et al. (2019), Keating (2007) and Ramsay (2001) concluded that páramo vegetation recovery does not follow one particular successional trajectory. Fires can have very different effects on vegetation, depending on the intensity and extent of a particular fire (Keating 2007; Luteyn 1999; Sklenář and Ramsay 2001; Suárez and Medina 2001; Zomer and Ramsay in review). Fire behaviour is determined by the pre-fire community structure (physical vegetation structure and fuel accumulation) and fire event conditions (Zomer and Ramsay in review).

Rates of tussock grass regeneration have been observed to be high (Gutiérrez-Salazar and Ramsay in press), but the impacts of fire on other páramo life forms vary widely (Horn 1989; Horn 1997; Janzen 1973; Keating 1998). Most páramo species have adaptations that help them to survive fires (Laegaard 1992), but the mechanisms vary among species, making the recognition of community-level successional trends at this taxonomic level difficult. A credible alternative for recognizing community assembly patterns is the use of plant functional types (Smith and Huston 1989): “nonphylogenetic groupings of species that show close similarities in their resource use and response to environmental and biotic controls” (Wilson 1999). To assist in such studies, Ramsay and Oxley (1997) proposed ten plant growth forms for the páramos, each summarising the responses of its component species.

One difficulty in developing a better understanding of páramo fire ecology is the lack of well-documented study sites. Some studies have observed post-fire recovery without knowing what conditions were like before or during the fire (e.g., Kovář 2001; Ramsay 2001). In other cases, fires have been set experimentally, sometimes with the addition of a fire stimulant (Keating 1998; Ramsay and Oxley 1996). Longer term studies are impeded by the lack of fire records, noting when, where and how páramo fires took place (Bremer et al. 2019; Horn and Kappelle 2009; Matson and Bart 2013).

Using a fire register for the Reserva Ecológica El Ángel, northern Ecuador, we recorded diversity and abundance of growth forms in a chronosequence (space for time substitution; Walker et al. 2010) of páramo sites burned from 2000–2014, and from a site that had not been burned for at least 40 years. We also measured light levels and soil temperatures in the same plots. Our aim was to determine the key phases in post-fire vegetation succession in the páramo, and the mechanisms driving the changes.

## Methods

### Study areas and fire records

The Reserva Ecológica El Ángel (REEA) is located in the Western Cordillera of northern Ecuador, and protects part of a contiguous páramo that is shared with Colombia. The buffer zone around the reserve is dominated by agricultural land use, where fires often occur. In the past, fires have also been common in parts of the reserve itself. Typically, such fires were started to stimulate livestock forage, aid hunting, or happened by accident (Ramsay 2001). An inventory of known fires since 2000 in the reserve and its buffer zone has been kept with the help of reserve officials and local fire brigades (Bustos Insuasti 2008; Valdospinos Navas 2008). More recently, the fire brigade in San Pedro de Huaca, Carchi, has also begun recording páramo fires for parts of the Eastern Cordillera in northern Ecuador, providing further opportunities for study.

Using these records, twelve sites in REEA and its buffer zone (Western Cordillera) and three additional sites from the páramo of La Bretaña (Eastern Cordillera) were selected for study, representing fires that burned from 2000–2014, from <1 to 15 y before our survey, at elevations of 3500–3900 m. Each fire site was located with GPS coordinates obtained from the records. In addition, one site known to be unburned for at least 40 years was included, making a total of 16 study sites. The páramo grasslands of these study areas were dominated by *Calamagrostis* tussock grasses and giant rosettes of *Espeletia pycnophylla* Cuatrec. (at densities of 1300–5400 adult plants ha^-1^).

### Data collection in the field

One 50m x 2m plot was randomly selected within each fire site. The plot was divided into 100 x 1 m^2^ quadrats. In each of these 1 m^2^ quadrats, the presence or absence of ten páramo plant growth forms, defined by Ramsay and Oxley (1997), was recorded: stem rosettes, basal rosettes, tussock grasses, acaulescent rosettes, cushions/mats, upright shrubs, prostrate shrubs, erect herbs, prostrate herbs, and trailing herbs.

Vegetation height was measured in each 1 m^2^ quadrat using the drop-disc technique (Stewart et al. 2001), with a 20 cm diameter disc, 190 g in weight, allowed to fall to rest on top of the vegetation (excluding *Espeletia*).

Soil temperature was measured at 20 cm depth using Signstek 3 1/2 6802 II Dual Channel Digital Thermometer with 2 K-type thermocouple sensor probes. Measurements were taken at five regular intervals along the longest axis of each plot, at 5 m, 15 m, 25 m, 35 m, and 45 m. At each place a temperature reading was taken in each of three shading conditions: beneath dense tussocks (shaded), on the edge of tussocks (intermediate) and in open intertussock areas (open).

Light at ground level was determined by measuring the percentage of incident photosynthetically active radiation (PAR) using a SunScan Canopy Analysis System with BF2 Beam Fraction Sensor (Delta-T Devices Ltd, Cambridge, UK). The sunscan probe was held at ground level underneath the vegetation at 1 m intervals along the length of the plot (50 in total). Each reading consisted of 64 simultaneous measurements of the percentage of total light (above the canopy) reaching the sensors on the ground. The median percentage of the light above the canopy which reached ground level was calculated for each 1 m interval along the transect, and subsequent analysis was based on these median values (*n*=50 for each plot).

Shannon’s diversity index (using log_e_) was calculated for growth forms for all sites. Non-metric multidimensional scaling (MDS) was carried out with Primer 6 (PRIMER-E, Plymouth, UK) to compare growth form composition between the fire intensity plots. Other standard statistical tests were performed with R version 3.4 (R Core Team 2019).

## Results

Diversity of growth forms in sites through time after fire followed a three phased pattern (Fig 1). In phase 1, there was a rapid increase to the highest levels of diversity within 2 years. Unfortunately, we had access to just one site <1.5 y after fire. In phase 2, diversity decreased, with the lowest levels occurring 8–10 y after fire. The bulk of our plots were burned within this time range. In phase 3, diversity increased from 10 to 15 y after fire and was even higher in the plot which had not been burned for at least 40 y, almost reaching the same level of diversity registered in at the start of phase 2.

**Fig. 1.**
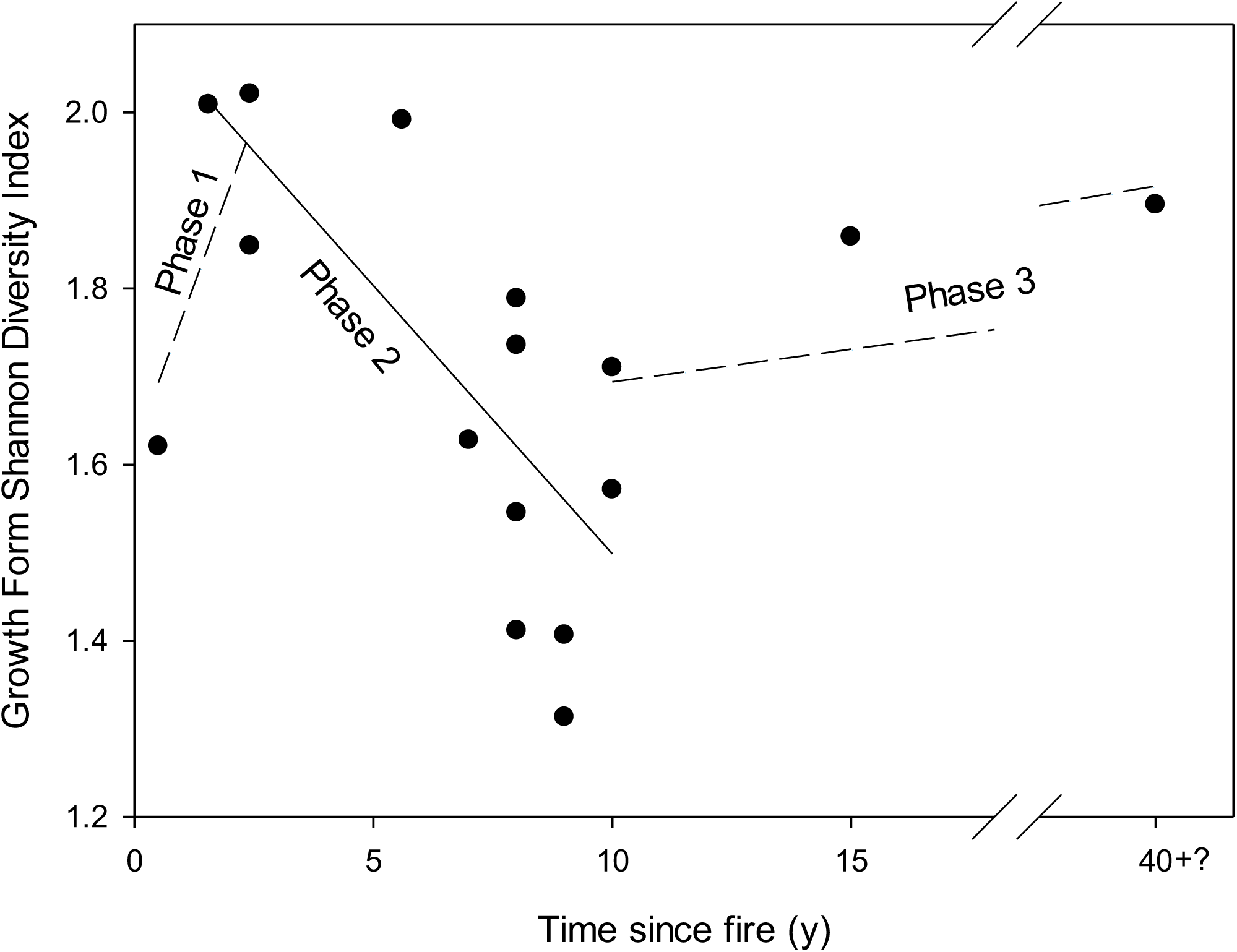
Three phases of plant growth form diversity response through time after fire for 16 plots. The plot at 40 y since fire was not burned during this time and might have been unburned for even longer. Part of the *x*-axis has been omitted.

Changes in the composition of plant growth forms did not follow a straightforward pattern (Fig. 2). Community compositions were fairly similar among sites 1.5–5.6 y after fire and among sites 8–10 y after fire. The unburned control was most similar in composition to sites 1.5–5.6 y after fire.

**Fig. 2.**
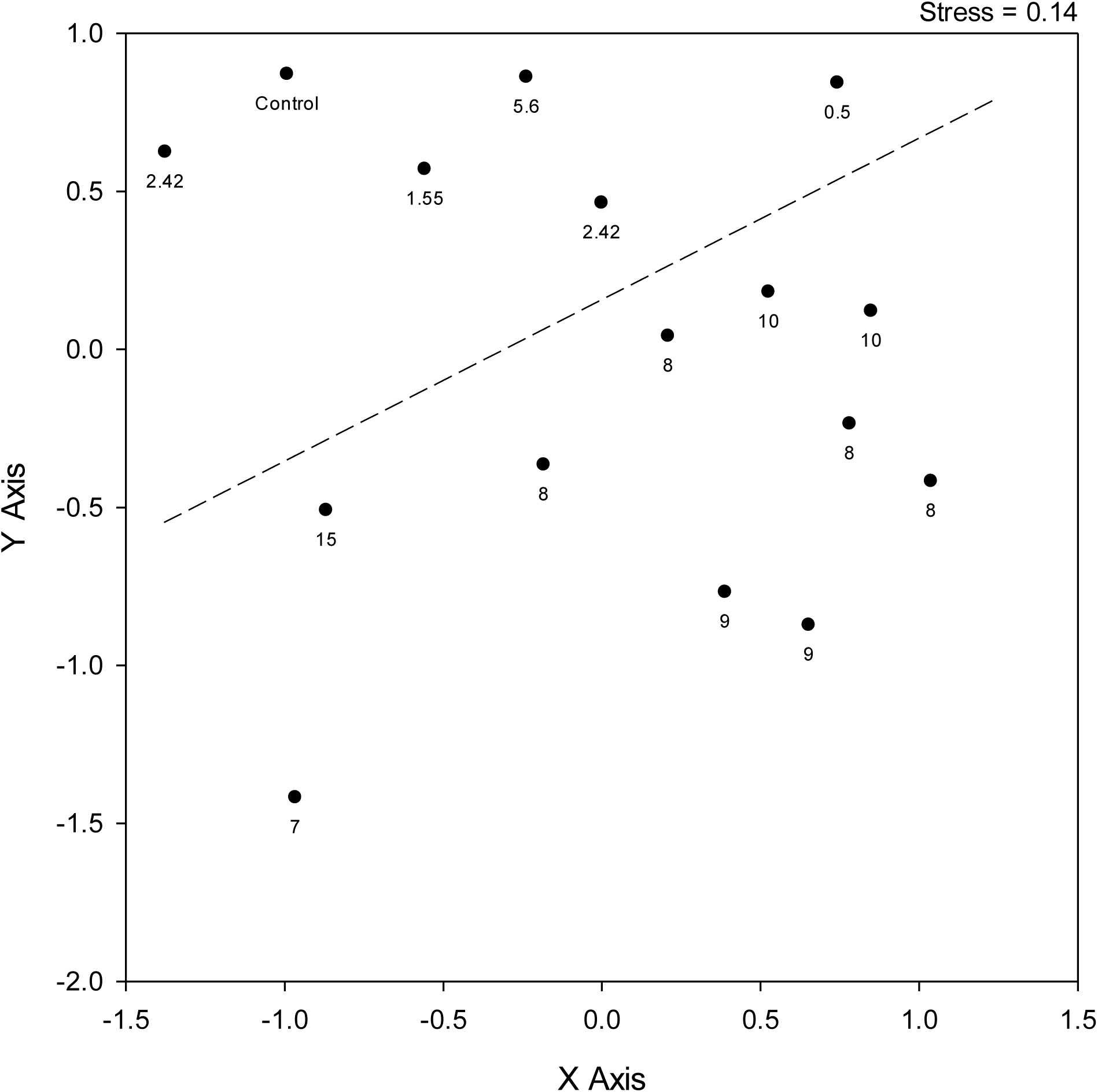
Non-metric Multidimensional Scaling ordination of plant growth form composition based on presence absence data for 100m^2^ plots. Each plot is represented by a point, labelled with the time since fire in years. The distance between plots in the ordination diagram shows the difference between them in growth form composition. The further a site is from another in the diagram, the greater the difference in composition. The dotted line highlights a divide between plant growth form assemblages 0.5–5.6 year since fire and 7–10 y since fire.

Analysis of only those sites burned 1.5–10 y after fire (Phase 2 in Fig. 1) shows that diversity decreased as time since fire increased (regression: *F*_1,11_=14.3, *p*=0.003; Fig 3a). Richness of growth forms did not vary significantly through time from 1.5 to 10 y after fire (regression: *F*_1,11_=0.145, *p*=0.710; Fig. 3b). The majority of sites had nine of the ten growth forms present. Lowest richness was found in two sites, both 9 y after fire (five and seven growth forms respectively).

**Fig. 3.**
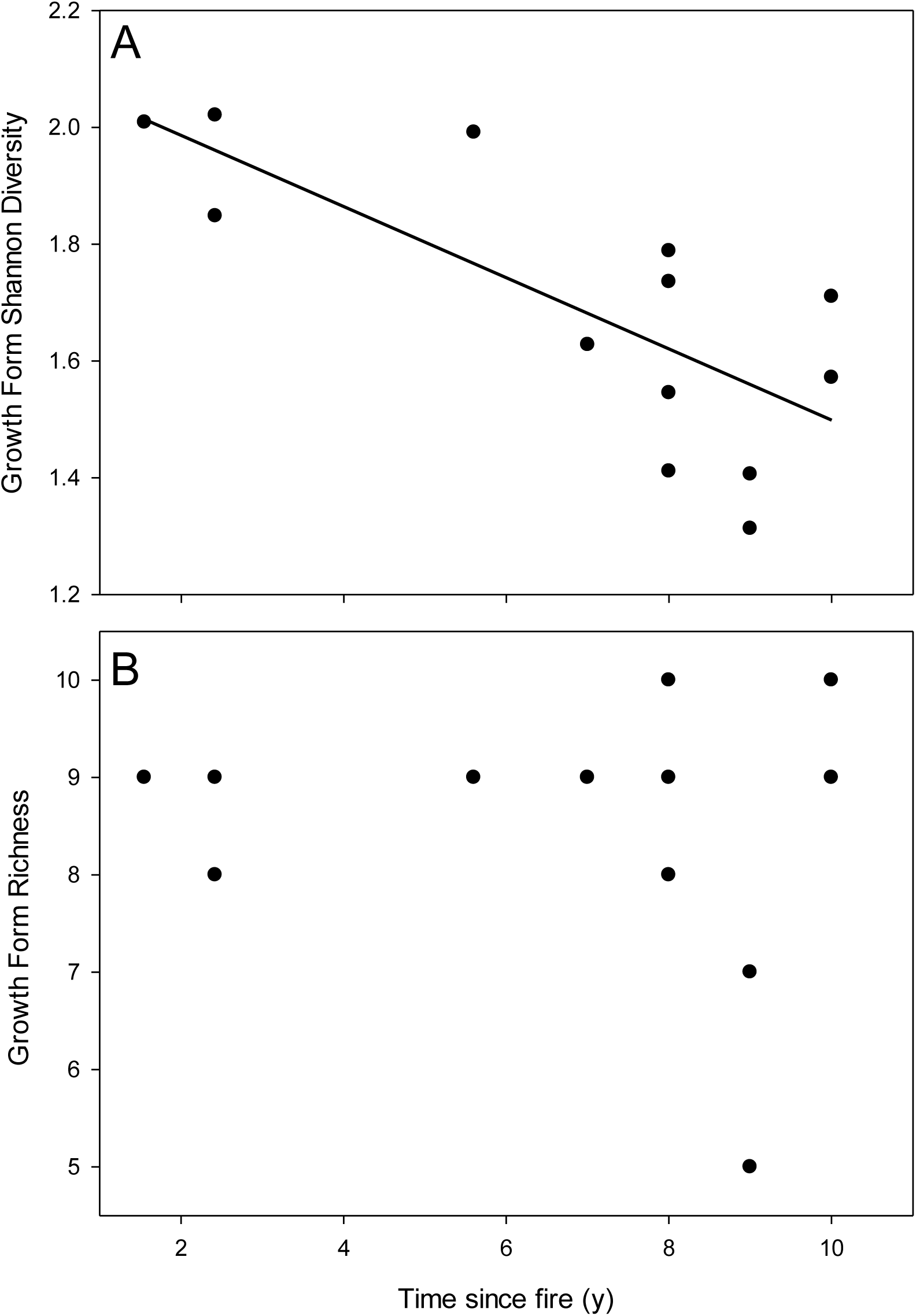
Growth form diversity and richness in thirteen páramo sites, 1.5–10 y after fire. A. Shannon’s diversity index (*R*^2^=0.565, fitted line *y=*2.11–0.06*x*). B. Growth form richness (of 10 growth forms).

The frequencies of several growth forms within the 100 m^2^ plots decreased as time since fire increased from 1.5 to 10 y: prostrate herbs (regression: *F*_1,11=_10.1, *p=*0.009; Fig. 4A), prostrate shrubs (regression: *F*_1,11=_17.9, *p=*0.001; Fig. 4B), cushions and mats (regression: *F*_1,11=_6.9, *p=*0.024; Fig. 4C), and giant basal rosettes (regression: *F*_1,11=_9.3, *p=*0.011; Fig. 4D).

**Fig. 4.**
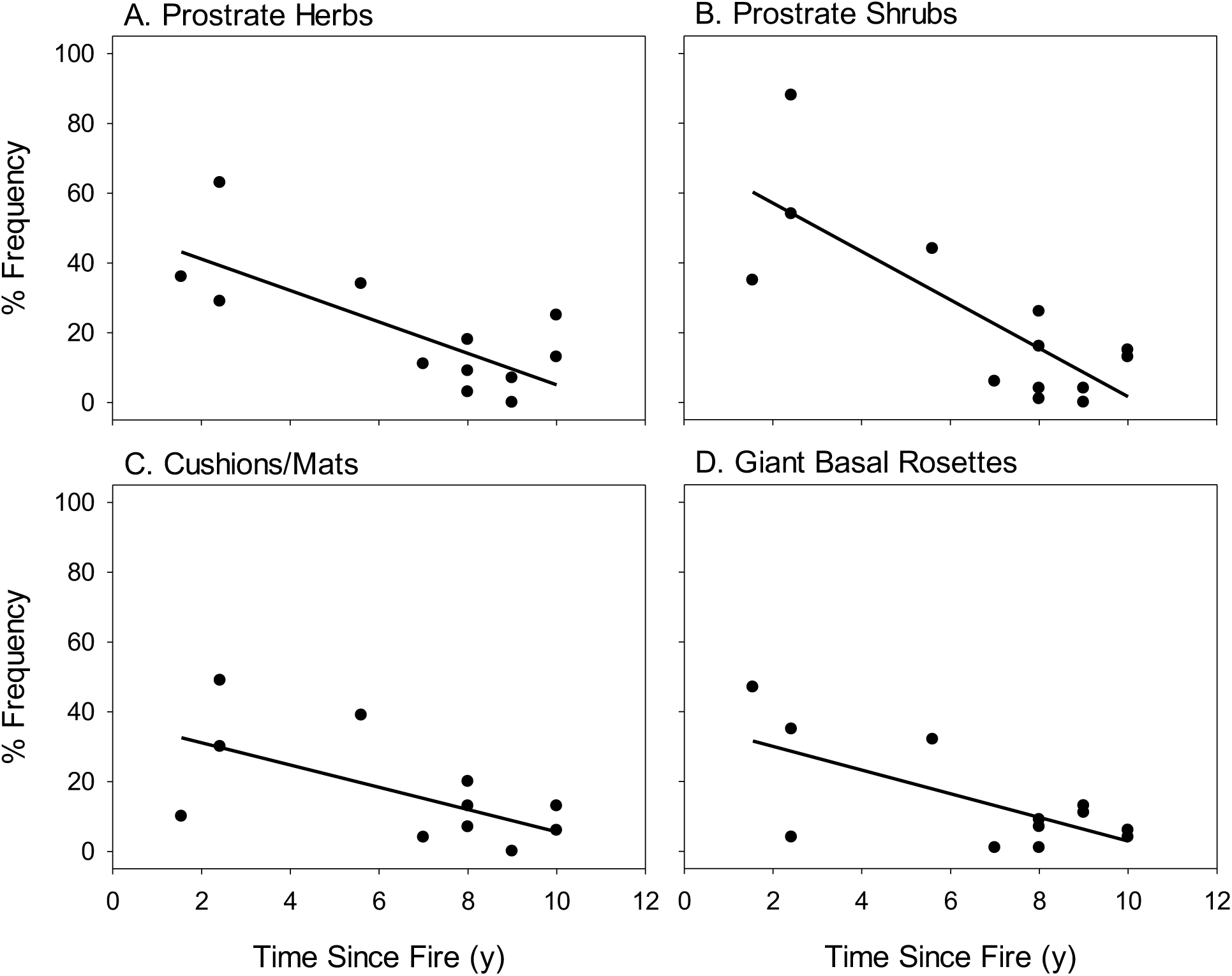
Frequency of four plant growth forms in 100m^2^ plots of differing times since fire, 1.5–10 y after fire. (a) prostrate herbs (*R*^*2*^= 0.551, *y*=50.14–4.5*x*) ; (b) prostrate shrubs (*R*^*2*^=0.621, *y*=71.01-6.93*x*) ; (c) cushions & mats (*R*^*2*^=0.385, *y*=37.52–3.19*x*); (d) giant basal rosettes (*R*^*2*^=0.459, *y*=36.82–3.39*x*).

Six growth forms did not follow a distinct pattern of changing frequencies from 1.5–10 y after fire. Tussock grasses were present at high frequency in all plots (99%–100%; Fig. 5A). Giant stem rosettes varied in frequency (40–99%; Fig. 5B), as did upright shrubs (20–65%; Fig. 5C) and erect herbs (1–81%; Fig. 5D). Acaulescent rosettes were found at low abundance (0–11%; Fig 5E), and trailing herbs were absent from most plots and were found just once each in two plots (Fig. 5F).

**Fig. 5.**
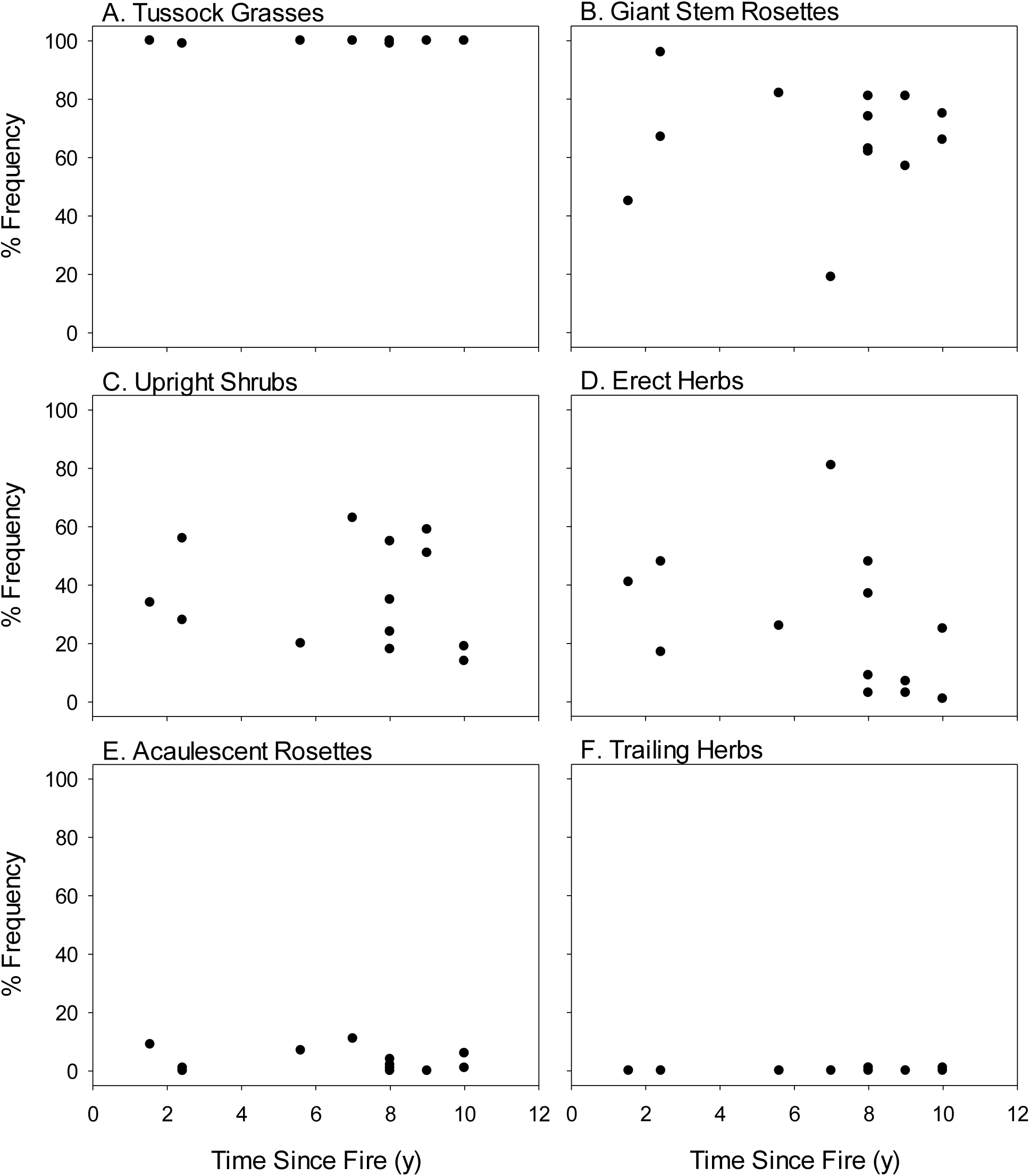
Frequencies of six growth forms in 100 m^2^ plots in plots 1.5–10 y after fire. A. Tussock grasses. B. Giant stem rosettes. C. Upright shrubs. D. Erect herbs. E. Acaulescent rosettes. F. Trailing herbs.

Mean vegetation height increased by approximately 25 cm on average as time since fire increased from 1.5–10 y (regression: *F*_1,11_=13.7, *p*=0.040; Fig 6A). In contrast, soil temperatures at depth decreased (regression, *F*_1,11_=34.3, *p*<0.001; Fig 6B). Median percentages of incident PAR reaching ground level decreased (regression: *F* _1,647_= 491.0, *p*<0.001,; Fig. 6C), while the proportion of ground with heavy shading (<10% incident light at ground level) increased as time since fire increased from 1.5–10 y since fire (regression: *F* _1,11_=23.7, *p*<.001; Fig. 6D). For context, the plot unburned for at least 40 y had a median PAR of 12.4% and the proportion of the readings with heavy shading was 62%.

**Fig. 6.**
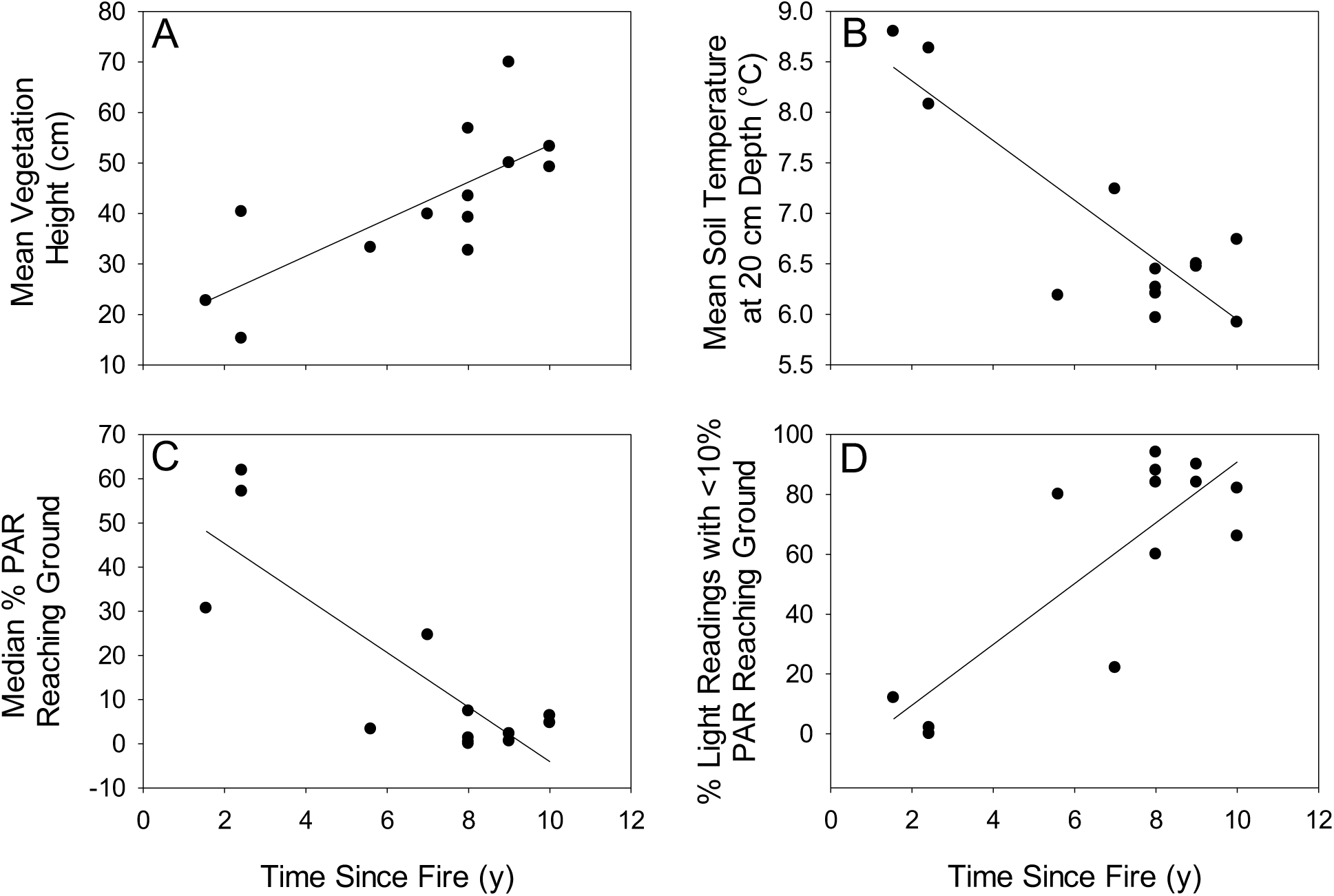
Changes in vegetation height, soil temperature and light conditions at the soil surface. A. Mean vegetation height (*R*^2^=0.553, *y*=16.91+3.67*x*). B. Mean soil temperatures at 20 cm depth (*R*^2^=0.757, *y*=8.9–0.3*x*). C. Medians of photosynthetically active radiation (PAR) reaching ground level, as a percentage of the incident light above the vegetation canopy (*n*=50 for each plot; *R*^2^=0.431, *y*=55.5–5.51*x*). D. Proportion of medians that were <10% incident PAR reaching ground level (*n*=50 for each plot; *R*^2^=0.683, *y* =10.16*x*–10.78).

## Discussion

This study considered broad patterns in plant growth forms, since it is the interaction between these forms that helps to explain the mechanisms of succession. However, many distinct species belong to each growth form, and do not all respond to fires in a similar way. Further investigation is needed at the species level to fully understand the implications for biodiversity of different successional trajectories or fire suppression. Particular attention should be focused on endemic species and other species of conservation concern (Matson and Bart 2013), as well as the contribution of different páramo growth forms to the provision of ecosystem services (Bremer et al. 2019).

Although community assembly dynamics are known to be highly variable after páramo fires (Keating 2007; Ramsay 2001), our study highlighted three key phases in terms of growth form diversity and composition. The mechanisms responsible for these successional patterns are proposed here to be the consequence of differential survival and recruitment after fire, followed by competitive interactions between growth forms, driving temporal changes in environmental conditions as the structure of vegetation developed (summarised in Fig. 7, and discussed in more detail below).

**Fig. 7.**
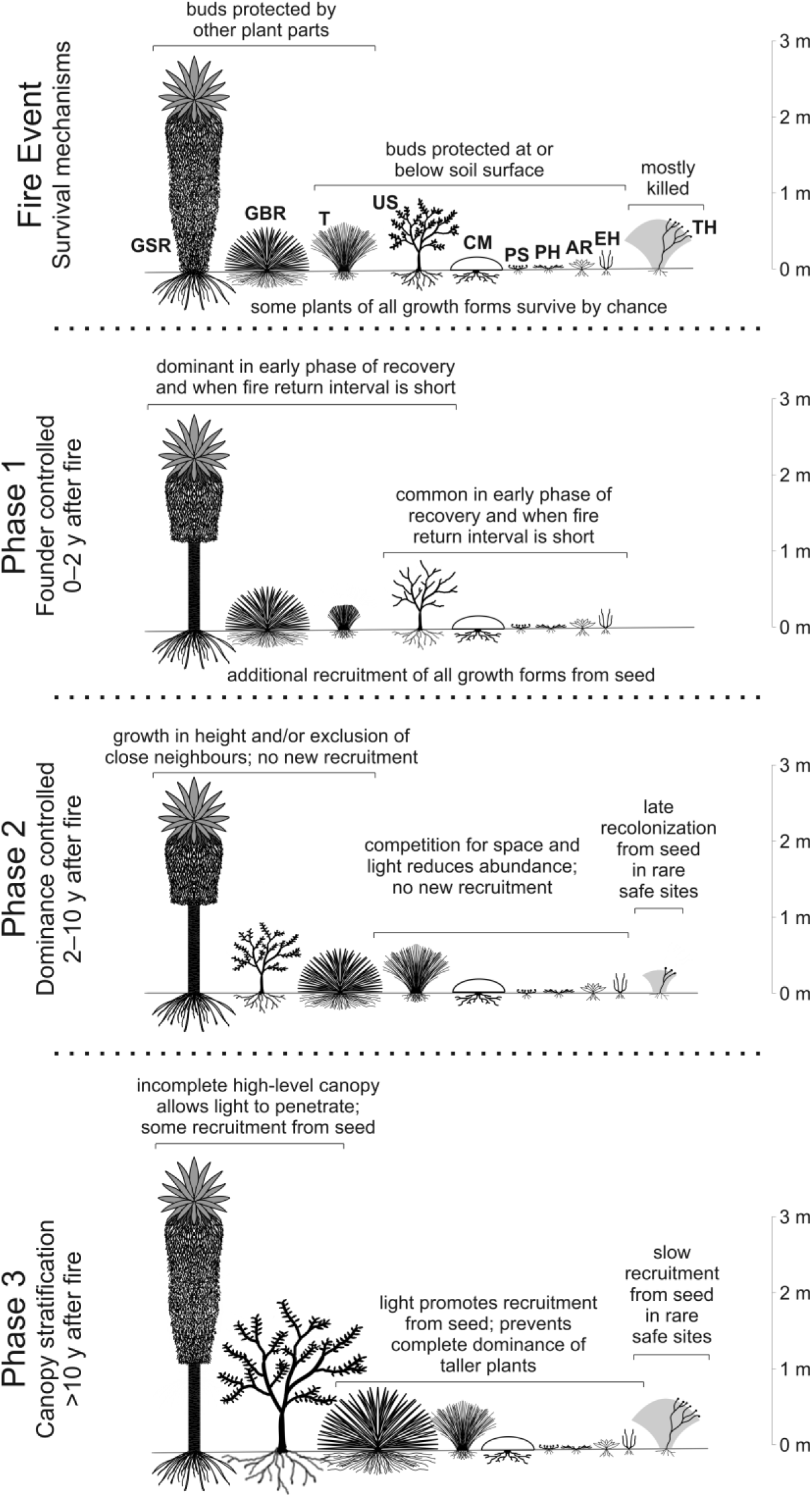
Schematic summary of responses of ten páramo growth forms to fire and their interactions during three phases of post-fire succession. Growth forms: GSR = giant stem rosette, GBR = giant basal rosette, T = tussock, US = upright shrub, CM = cushion or mat, PS = prostrate shrub, PH = prostrate herb, AR = acaulescent rosette, EH = erect herb, TH = trailing herb.

### Fire event leading to Phase 1

Vegetation composition soon after a fire is determined by plants’ fire survival strategies and their regeneration mechanisms immediately after fire. Subsequently, most plants are able to grow well in light conditions and warmer soil temperatures. This phase can be considered founder-controlled, where the abundance of plants is driven by survival and recruitment processes.

Tussock plants are rarely killed by fires (Hofstede et al. 1995) as their deeply-seated shoot apices and buds are protected against heat in the centre of the tussock base (Laegaard 1992; Ramsay and Oxley 1996). Nevertheless, partial mortality is often observed, leading to fragmentation of plants and a smaller basal area (Gutiérrez-Salazar and Ramsay in press; Laegaard 1992; Ramsay 1999; Suárez and Medina 2001). In the initial recovery phase after a fire, there is a tussock-building phase caused by the growth of surviving fragments and recruitment of new seedlings in open areas (Ramsay 2001; Ramsay and Oxley 1996; Vargas-Ríos 1997; Vargas Ríos 2011). During this period, the height of tussocks is restricted, permitting more light at ground level and increased soil temperatures (Gutiérrez-Salazar and Ramsay in press). This allows other growth forms time and space to recover after the fire.

Adult giant stem rosettes tolerate fire well, protecting apical buds in at the heart of their rosettes (Laegaard 1992). Germination and establishment of *Espeletia pycnophylla*, co-dominant in our study plots, is known to be enhanced after fires (Laegaard 1992; Suárez and Medina 2001). This is consistent with the initial rise in frequency of giant stem rosettes in our study, which also included *Blechnum* stem rosette ferns.

Giant basal rosettes (*Puya hamata* in our plots) also survive fires well by shielding their buds against high fire temperatures (Garcia-Meneses and Ramsay 2014). Recruitment of new *Puya* plants can be enhanced significantly in the open conditions that are typical after a páramo fire, but only if seed-producing adults are present nearby since seed dispersal is poor (Garcia-Meneses and Ramsay 2014; Miller and Silander 1991). This can result patchy distribution patterns of *Puya* at the landscape scale, with consequences for the behaviour of pollinating hummingbirds and the genetic diversity of the plants and their seeds (Garcia-Meneses and Ramsay 2012; Rivadeneira et al. 2020).

Some upright shrubs regenerate well after fire, from insulated buds or roots (Horn and Kappelle 2009; Ramsay and Oxley 1996) and the abundance of some species might benefit from repeated burning (Keating 2007). The abundance of upright shrubs in our plots did not change markedly with time since fire, suggesting plants maintained their presence from the early stages of recovery, perhaps with occasional colonization from seed.

Other growth forms survive fires at or below the soil surface, where fire temperatures are lower (Ramsay and Oxley 1996): cushion and mats, acaulescent rosettes, prostrate herbs, prostrate shrubs, and erect herbs. Survivors of these growth forms, plus new plants recruited from seed rain (Vargas-Ríos 1997), benefit from the open conditions during this initial phase to establish and grow.

Trailing herbs, usually growing through the canopies of tussocks do not survive fires well, since they hold their buds in the zone of maximum fires temperatures (Ramsay and Oxley 1996). Plants belonging to this growth form are less abundant in burned páramos, and they were completely absent from our plots in the early phase of post-fire recovery. It is worth noting that even growth forms which are vulnerable to mortality from fire can survive by chance in small islands of vegetation that the fire missed (Laegaard 1992). The relative abundance of these islands, and their later importance in determining the composition of the vegetation merits more attention.

### Phase 2

Phase 2 of recovery showed a decrease in growth form diversity from 1.5–10 y after fire, though growth form richness did not vary greatly. The compositional analysis also supports the conclusion that the changes reflect shifts in relative abundance rather than local exclusion of growth forms.

In this second phase, tussock grasses increase their dominance. Gutiérrez-Salazar and Ramsay (in press) have shown that the height of the tussocks increases steadily during this phase, resulting in a consistent shading effect and decline in temperature at ground level. The dominance-controlled plant community during Phase 2 is thus characterized by the suppression of potential competitors by the tussocks.

The lack of light and lower temperatures at the soil surface inhibits the establishment of growth forms that grow over the soil surface (prostrate herbs, prostrate shrubs, cushions). Although large giant basal rosettes prevent neighbouring tussocks from shading their leaves, smaller plants are vulnerable to being outcompeted for light by taller tussocks. It is this loss of smaller plants and low recruitment of new plants beneath the tussock canopy that explains their decline during Phase 2 (Garcia-Meneses and Ramsay 2014).

Long-lived giant stem rosettes and upright shrubs did not show any consistent trend in response to time since fire during this second phase of succession. Their height means they are mostly unaffected by the density of the tussock grass canopy.

The abundance of acaulescent rosettes was relatively low, but stable, throughout this phase. These plants favour the canopy gaps where more light reached the ground (all plots had some places which received 50–60% incident light at ground level). Although these gaps were rare in plots representing the later stages of phase 2, they still provided appropriate conditions for some acaulescent rosettes. Erect herbs often grow alongside the tussock leaves in the canopy, or in canopy gaps, and did not show a trend through time in Phase 2.

Trailing herbs were only found in sites 8–10 y after fire. They rely on physical support of tussock grasses, but their late arrival suggests poor seed dispersal and/or germination for the species present in our study area.

It is noteworthy that one plot, 7 y after fire, was dissimilar in composition to the other plots. It was located on the edge of REEA’s buffer zone, adjacent to agricultural fields, and was the only plot in this study to be regularly grazed by cows. This grazing had a visible effect on the vegetation with large spaces between tussocks, providing suitable light conditions for a greater abundance of erect herbs and acaulescent rosettes, but with far fewer *Espeletia* stem rosettes. Grazing disturbance often occurs alongside fire in páramo grasslands and such combined disturbance regimes maintain fragmented tussock grasses and more open conditions (Hofstede et al. 1995; Ramsay and Oxley 1996; Verweij and Kok 1992).

### Phase 3

The longer-term absence of fire in some parts of our study area gave some insight into successional change in plots 10 y or more after fire: Phase 3 in our proposed scheme. During this phase, the diversity of growth forms increased with another shift in relative abundances. Our “control” plot, unburned for at least 40 y, had growth form diversity and composition similar to the highest levels recorded in sites 1.5–5.6 y after fire. This was an unexpected trajectory, and while it is important to be cautious before assuming this observation is typical, it raises some interesting ideas about what might happen if more páramo areas were fire-free.

In our study, the 15 y since fire and the older “control” site had notably less tussock grass cover (only 89% and 79% frequency respectively) and the highest frequencies of upright shrubs (68% and 73% respectively). It seems that upright shrubs begin to outcompete the tussock grasses as time since fire passes. Such woody encroachment into unburned grassy páramo has been suggested before (e.g., Laegaard 1992), and could be maintained by positive feedbacks in temperature, soil moisture or nutrient availability (Brandt et al. 2013; Matson and Bart 2013). Woody encroachment is a concern to environmental managers of grasslands because it can alter ecosystem structure and function (Knapp et al. 2008; Zavaleta and Kettley 2006) and can lead to declines in biodiversity (Costello et al. 2000; Ratajczak et al. 2012). The implications of growth form shift in unburned páramos for services such as water provision and carbon storage is unclear. High vegetation cover of tussock grasses is often associated with protecting and promoting the ecosystem function of páramo soils, *i.e.*, providing water regulation, storing and sequestering soil carbon (Bremer et al. 2019; Minaya Maldonado 2017). Molina et al. (2019) demonstrated higher rates of chemical weathering in páramo-zone soil under trees with soil beneath tussock grassland in southern Ecuador.

While our study and those of Matson and Bart (2013) and Bremer et al. (2019) suggest that diversity and growth form richness may not decrease in a shrub dominated páramo, species composition is likely to shift. It is not clear how many páramo species require disturbance gaps for their survival, but it is likely that many species of conservation interest would see reductions in their abundance in areas where burning was prevented for many decades.

From an ecological perspective, the existence of páramo without fire is a relatively recent and rare phenomena (White 2013) and is only beginning to be studied. If fire suppression does lead to the transition from grass páramo to a shrub dominated páramo with potentially altered ecosystem function, the potential consequences are of conservation concern and should be evaluated (Armenteras et al. 2020; Matson and Bart 2013). It is important to recognise that conservation-motivated policies to exclude fire in páramo areas might result in different ecological outcomes from those desired by the policy makers.

## Conclusions

We suggest that the recovery of páramo vegetation after fire comprises three phases. Immediately after fire, a survival and recruitment phase occurs in more open conditions, with high diversity of growth forms. As time passes, the growth of tussock grasses prevents many other plants from establishing, reducing diversity. With yet more time, shading from taller plants thins out the tussock cover nearer the ground, allowing certain growth forms to establish which had been mostly excluded in the previous phase—leading to a return of growth form diversity. We have less confidence in this last phase, because of a lack of replication in our study, but it is consistent with observations reported by other researchers.

At present, the prominent fire management strategy in many páramo regions is to prohibit fires, with the aim of protecting the integrity of the grassland ecosystem, promoting carbon storage and water provision. It is not yet clear whether this strategy will result in the desired outcomes. Some authors have concluded that this conservation strategy is unrealistic and difficult to enforce with local farmers (Keating 2007), that total burn exclusion is unnecessary to conserve plant species richness, growth form richness or vegetation cover (Bremer et al, 2019), and that it is not consistent with the environmental and cultural history of these fire-dependent páramos (Horn and Kappelle 2009; White 2013). We agree with this assessment and urge policy makers to consider the ecological evidence associated with the strategies being adopted. At very least, long-term studies should be employed to monitor future changes in composition and ecosystem function in response to policies of fire suppression. It would be prudent to support important policy decisions with careful evidence-based consideration of the outcomes.

## Acknowledgements

This work was carried out as part of permit MAE-DPAC-UPN-BD-IC-FLO-2015-004, issued by the Ecuadorian Ministry of Environment. Fieldwork was carried out by the authors, with assistance from Anna Masters, Cheryl McAndrew, Patricia Gutierrez Salazar, Alejandro Marchán & Juan Yépez Cardenas. Logistical support in REEA was provided by the reserve’s administration and rangers, who also provided information from fire records. The fire brigade in San Pedro de Huaca gave us fire dates for the sites at La Bretaña.

